# *Cassava common mosaic virus* causes photosynthetic alterations associated with changes in chloroplast ultrastructure and carbohydrate metabolism of cassava plants

**DOI:** 10.1101/2020.04.23.057604

**Authors:** Andrea A. Zanini, Liliana Di Feo, Dario F. Luna, Pablo Paccioretti, Agostina Collavino, Marianela S. Rodriguez

## Abstract

*Cassava common mosaic virus* (CsCMV) is a potexvirus that causes systemic infections in cassava plants, leading to chlorotic mosaic and producing significant yield losses. To date, the physiological alterations and the mechanism underlying biotic stress during the cassava-CsCMV compatible interaction remains unknown. In this study, we found that CsCMV infection adversely modified chloroplast structure and had functional effects on chloroplasts in source leaves during the course of viral infection. Extrusion of the chloroplast membrane with amoeboid-shaped appearance was observed in infected mesophyll cells. These alterations were associated with lower relative chlorophyll content, and reduced PSII efficiency and CO_2_ fixation. Moreover, an oxidative stress process was observed in CsCMV-infected plants. Strong declines in the maximum quantum yield of primary photochemistry (F_v_/F_m_) were observed in infected plants. Furthermore, the analysis of Chlorophyll-a fluorescence (ChlF) evidenced a progressive loss of both oxygen evolving complex activity and “connectivity” within the tripartite system (core antenna-LHCII-Reaction Centre). Other effects of the pathogen included reduction of starch and maltose content in source leaves, and a significant increase of the sucrose/starch ratio, which indicates alteration pattern of carbon. Our results suggest that CsCMV induces chloroplast distortion associated with progressive chloroplast function loss and diversion of carbon flux in source leaf tissue, which should be key in inducing yield losses of infected crops.

**Main conclusion:** CsCMV infection adversely modified chloroplast structure and had functional effects on chloroplasts during the course of viral infection, associated with metabolic adjustment in cassava source leaves, which would partly explain cassava root yield losses.

## Introduction

Cassava (*Manihot esculenta* Crantz) is one of the most widely grown tuber crops and an important food security crop for smallholder farmers, especially in low-income, food-deficit areas (Bellotti et al. 1999). Its storage roots are the main organ, which is marketed as feed resource and feedstock for starch extraction, bio-energy production, and pharmaceutical and textile industries; accordingly, cassava is a widely recognized sustainable crop in the context of the global changing climate (FAO 2013; De Souza et al. 2017; Marx 2019). Therefore, its productivity is conditioned by the starch accumulation from CO_2_ fixation and the source-sink relationship (Yan et al. 2019). Since cassava is vegetatively propagated, its yield is greatly restrained by viral diseases (Legg et al. 2015). Among all reported cassava-infecting potexviruses (Lozano et al. 2017), *Cassava common mosaic virus* (CsCMV) has been identified as the only one able to cause disease in single infection and to produce significant yield losses (Costa and Kitajima 1972; Venturini et al. 2016; Zanini et al. 2020). This pathogen has been detected infecting cassava crops in South America (Calvert et al. 2012; Chaparro-Martinez and Trujillo-Pinto 2003; Silva et al. 2011; Fernandez et al. 2017; Zanini et al. 2018) and recently in China (Tuo et al. 2020).

The typical symptoms of CsCMV infection, known as cassava common mosaic disease (CCMD), vary in severity and include a wide range of foliar chlorotic mosaic (light green chlorotic areas, interspersed with dark green areas), sparsely distributed in the leaf blade of cassava plants (Zanini et al. 2018). This kind of symptoms might be the consequence of chloroplast deformation and shut-off induced by the virus infection (Li et al. 2016a). The viral influence on chloroplast structure and function, which usually leads to photosynthetic impairment, is common across diverse plant-virus interactions (Zhao et al. 2016). Accordingly, viruses interfere with photochemical reactions by reducing the abundance of key proteins involved in the photosynthetic electron transport chain (Souza et al. 2019; and references therein). In this context, alterations in electron transport might generate reactive oxygen species (ROS) (Rodríguez et al. 2010). The ROS-antioxidant balance determines, to a large extent, the cellular redox state, which plays a central role in environmental stress perception and defense response modulation (Foyer and Noctor, 2005).

To enable their multiplication, plant viruses have the ability to rearrange host metabolism and cellular machinery (Souza and Carvalho 2019). A wide range of physiological and biochemical disorders are triggered from cell infection, including relocation of photoassimilates, redox imbalance and premature senescence, with significant economic losses (Rodríguez et al. 2010, 2012; Souza and Carvalho 2019). Viral infections lead to a decline in CO_2_ fixation, which could be directly related to a decrease in carbohydrate accumulation, reducing plant growth and development (Sade et al. 2013; Nuwamanya et al. 2017). On the contrary, other studies have demonstrated that soluble sugars and starch accumulate in the infected leaves, where photosynthesis is reduced (Rodriguez et al. 2010; Andreola et al. 2019; De Haro et al. 2019). Furthermore, abnormal accumulation or depletion of starch in systemically infected tissues was reported in *Cucumber mosaic virus* (CMV, Cucumovirus)-marrow interaction (Técsi et al. 1996). Therefore, the mechanisms associated with changes in carbohydrate metabolism are specific to the plant-virus pathosystem; to our knowledge, the effect of CsCMV infection on sugar and starch content has still not been studied.

Given the significant yield losses caused by CsCMV (from 30% to 60%) (Costa and Kitajima 1972; Venturini et al. 2016; Zanini et al. 2020), it is imperative to investigate the physiological basis underlying cassava-CsCMV interplay. To extend the knowledge acquired in previous studies by Tascon et al. (1975), we focused our research on morphological and functional alteration of cassava chloroplast. Taking into account the theoretical background mentioned above, we hypothesized that CsCMV would induce anomalies in the chloroplasts associated with reduced photosystem II (PSII) electron transport activity and redox imbalance. We further proposed that since CO_2_ fixation decreases, sugar and starch content will be negatively affected. Therefore, the present study provides, for the first time, information about chloroplast structure and progression of the CCMD on chloroplast function in the compatible cassava-CsCMV interaction.

## Materials and Methods

### Plant materials

Twenty *in-vitro* grown, healthy cassava v. ICA Negrita (CM3306-4) plants were used. Sixty days after acclimation, 10 plants were graft-inoculated with budwood from CsCMV-affected cassava plants and the other 10 were autografted to maintain their healthy condition. Yellowing and blotchy mosaic symptoms were visible 20 days after inoculation. These plants were transplanted in the field for macro-propagation and as a source of stem cuttings for subsequent greenhouse trials.

### Experimental setup

Cassava plants were propagated clonally from segments (approximately 20 cm in length) of parental stems with at least 6 nodes. The plants were grown individually in plastic pots (3 L) containing a 3:1 mixture (v/v) of potting soil and sand mixture, in a naturally illuminated greenhouse. All experiments were carried out in summer (average temperature 28 ± 5 ºC). Thirty cassava stakes were used for each sanitary condition: non-infected plants (control) and CsCMV-infected ones, arranged in three completely randomized blocks.

Measurements of number of nodes and number of leaves per bud, SPAD and chlorophyll fluorescence were taken in three ontological times: 60, 75 and 90 days after planting (DAP) of cassava plants, encompassing the key stages of cassava growth, from the ‘seedling’ stage (young cassava plants that start from stem cuttings, until 60 DAP) to the formation of root system (until 110 DAP). Height was measured at 60, 75, 90 and 210 DAP (cassava growth stage of starch maturity) and photosynthetic rate was estimated at 90 and 210 DAP.

Leaf tissue samples (excluding petioles) were taken from the fourth leaf (counting from the base to the top) of each plant at 90 DAP. They were immediately frozen in liquid nitrogen and stored at −80 °C for subsequent biochemistry determinations. These leaves represent a source organ.

### Serological virus detection

CsCMV was detected by double antibody sandwich enzyme-linked immunosorbent assay (DAS-ELISA) performed in polystyrene microtitre plates with anti-CsCMV-IgG as described Zanini et al. (2018). All plants were tested.

### Transmission electron microscopy - Immunogold labelling

Small tissue sections (0.2 cm wide × 0.5 cm long) sliced from the middle part of CsCMV-infected leaves were fixed in 2% paraformaldehyde and 2% glutaraldehyde in 0.01 M phosphate buffer, pH 7.2, for 1 day. Then, they were postfixed in 1% OsO_4_, in the same buffer and embedded in LRGold, as previously described (Kitajima 1997, Maunsbach 1998). Sections were cut with a diamond knife using an ultra-microtome and collected onto nickel grids. For immuno-electron microscopy, sections mounted on nickel grids were preincubated with 1% bovine serum albumin (BSA) in 1% phosphate buffered saline (PBS) overnight, and then treated with specific antiserum anti-CsCMV (Zanini et al. 2018) for six hours. The sections were then exposed to gold conjugated protein A of 5 nm in diameter (Zymed, now part of the Invitrogen). Sections from uninfected cassava leaves were used as additional controls in the immunolabelling experiments. They were examined under a JEOL JEM-1200 transmission electron microscope (TEM, JEM 1200 EXII, Jeol, Tokyo, Japan).

### Chlorophyll content, chlorophyll-a fluorescence and gas-exchange measurements

#### Chlorophyll content

The relative chlorophyll (Chl) content was determined with a portable meter (Hansatech, Chlorophyll Content Meter), using dual wavelength optical absorbance measurements (620 and 940 nm). The measurements were taken simultaneously with Chlorophyll-a fluorescence (ChlF), as described below.

#### Chlorophyll-a fluorescence transient

ChlF emission was measured with a Pocket-PEA fluorometer (Plant Efficiency Analyzer, Hansatech Instruments Ltd., King’s Lynn, Norfolk, UK). The polyphasic OJIP fluorescence kinetics was used to evaluate the PSII activity in CsCMV-infected and non-infected cassava plants. Before ChlF measurement was taken, the fourth developed leaves were dark-adapted using leaf clips for at least 30 min in order to allow full oxidation of the reaction centers (RC); the minimum fluorescence or F_o_ or O step was obtained at ~ 50 μs. Then, an actinic 1- second light pulse of 3500 μmol photons m^−2^s^−1^ was applied to reach the maximum fluorescence emission (F_m_ or P step), at about 300 ms. The intermediate steps, called J and I, were recorded at 2 ms and 30 ms, respectively. The OJIP parameters were calculated using the Pocket-PEA manufacture software, following Strasser et al. (2004). The parameters are described in Supplementary Table S1.

Changes in the oxygen evolving complex (OEC) activity and in the connectivity at PSII were calculated according to Chen et al. (2016). To reveal changes in OEC (K-band), fluorescence curves were normalized at both the O and J steps, rendering W_OJ_ curves. Next, W_OJ_ curves from control plants were subtracted from treated ones (W_OJ(Treated)_ - W_OJ(Control)_) to disclose the K bands (ΔW_OJ_). Positive values in these bands denote damage to the OEC. To detect L bands, curves were normalized between 0 and 300 μs, and plotted as W_OK_ curves. The subtraction between W_OK(Treated)_ - W_OK(Control)_ was plotted to obtain ΔW_OJ_ curves, which allowed us to visualize L bands. The positive values of ΔW_OJ_ curves are proportional to the loss of connectivity at PSII.

#### Net photosynthesis rates

Net CO_2_ assimilation rates were measured in the fourth fully expanded leaves of cassava plants at 90 and 210 DAP with a portable photosynthesis system LI-6400 XT equipped with a LED leaf cuvette (Li-Cor, Lincoln, NE, USA). The concentration of CO_2_ inside the chamber was kept at 400 μmol CO_2_ mol^−1^. The light source was set at saturating incident photosynthetic photon flux density (PPFD) of 1500 μmol m^−2^s^−1^ (90% of red and 10% of blue light) and the temperature was maintained at 25 °C.

### Determination of the effect of CsCMV on cassava carbohydrate

#### Ethanolic leaf extract

Frozen tissue (75 mg) was ground with liquid nitrogen and cold homogenized with 750 μL of 80% ethanol. The extract was poured into a previously cooled microtube and centrifuged at 12,000 g at 4 °C for 10 min. The samples were kept on ice and aliquots were separated to measure the total antioxidant capacity using FRAP (*Ferric Reducing Ability of Plasma*). The remaining extract was resuspended in vortex and taken to a thermal bath at 80 °C for 20 min. The samples were then centrifuged at 12,000 g at 4 °C for 10 min and aliquots were separated from the supernatant to measure total sugars. The insoluble fraction was used for quantification of starch content.

#### Total soluble sugars

Quantification of total soluble sugars was adjusted to a final reaction of 200 μL in polystyrene plates. Ethanolic leaf extract and Anthrone reagent (1 μL each) were placed in each well (Fales 1951). The blank contained H_2_O and Anthrone reagent. The mixture was incubated at 4 °C for10 min, then at 80 °C for 30 min, and subsequently allowed to cool 20 min at room temperature. Glucose was used as standard to measure the total soluble sugars content, expressed as mg of glucose per g of FW.

#### Sugar determination (HPLC)

Frozen leaf samples (0.1 g FW) were ground to a fine powder with liquid nitrogen, homogenized and suspended in 0.9 mL hot 80% ethanol, and kept at 80 °C for 2 h. Then, 0.9 mL dH_2_O was added to each sample; samples were incubated at 99 °C for 10 min and centrifuged at 12,000 g for 5 min. The supernatant was filtered using a nitrocellulose filtration membrane (0.22 μm pore size). Sucrose, glucose, fructose and maltose were determined by high-performance liquid chromatography (HPLC) (Shimadzu) using an amine column, isocratic acetonitrile: water (81:19) flow (1 mL/min), at 30 °C. Sugars were identified by their retention times and quantified according to standards.

#### Starch content

Starch content was determined in the pellet, from reducing sugars released after hydrolysis with the enzyme α-amyloglucosidase (Sumner and Somers 1944), using glucose as a standard. Starch content in the source organs was expressed as mg per g of FW of plant material.

#### AGPase activity

The assay was performed in the reverse direction using phosphoglucomutase and Glucose-6-phosphate dehydrogenase to couple Glucose-1- phosphate formation in real time to NADP^+^ reduction and then measured by spectrophotometers. The crude extracts were obtained as described by Li et al. (2016b), adapted to cassava leaves. Briefly, 0.1 g of fresh tissue fully homogenized on ice in a 0.3 ml pre-cooled solution composed of 100 mM Hepes-NaOH at pH 7.4, 8 mM MgCl_2_, 2 mM EDTA, 12.5% (v/v) glycerol, 5% (v/v) polyvinylpyrrolidone, and 50 mM β - mercaptoethanol. Then, a 20 μL aliquot of the crude enzyme extract was used to start the assay as detail Li et al. (2016b). Absorbance values at 340 nm of the reaction solution were normalized by absorbance values at 340 nm resulting from the control. The AGPase activity was then estimated with the normalized absorbance values and was expressed in enzyme units per mg of protein, calculated according to Kulichikhin et al. (2016). Protein content was determined in leaf extracts by Bradford (1976) assay.

### Determination of Total antioxidant capacity (FRAP)

The FRAP assay was used to determine the total antioxidant capacity of the samples (Benzie and Strain 1996); the protocol was adjusted for performance in a final reaction of 200 μL in polystyrene plates. In each well, 2 μL of ethanolic extract solution (see ethanolic leaf extract) were placed to react with FRAP reagent in the dark at room temperature for 20 min and the absorbance at 600 nm was recorded. Trolox® (6- hydroxy-2,5,7,8-tetramethylchroman-2-carboxylic acid, Sigma-Aldrich) solutions of known concentration, within in the range of 12.5 and 87.5 μM, were used for calibration. Total non-enzymatic antioxidant activity was expressed as μmoles per g of FW.

### Statistical analysis

Experiments were performed in a completely randomized design. Mixed linear model was performed using InfoStat (Di Rienzo et al. 2019). Mean separation was accomplished using Least Significant Differences (LSD) via Fisher test at the 95% confidence level.

## Results

### Chlorotic symptom during CsCMV infection: effect on chloroplast ultrastructure

Cassava plants infected with CsCMV exhibited the typical symptoms of CCMD, such as foliar chlorotic mosaic and mild leaf distortion (Fig. 1). The mosaic symptom was evident since leaf appearance. Furthermore, those plants that were self-grafted remained asymptomatic (non-infected plants). Systemic spreading of virus in plant tissues was confirmed by DAS ELISA (data not shown). Even though viral infection showed conspicuous foliar mosaic, no differences in the fully expanded leaf area were observed between healthy and CsCMV-infected plants (Supplementary Fig. S1). Yet, growth analysis showed that viral infection significantly affected (p< 0.05) the absolute and relative elongation rate (AER and RER, respectively). Indeed, the analysis of AER of infected plants showed a rapid stimulation of plant height, with a subsequent decrease of the rate over the days with respect to non-infected plants (Supplementary Fig. S2). The number of nodes and leaves per node increased over time; however, no significant differences were observed between plants of different sanitary conditions (Supplementary Fig. S3).

**Fig. 1.**
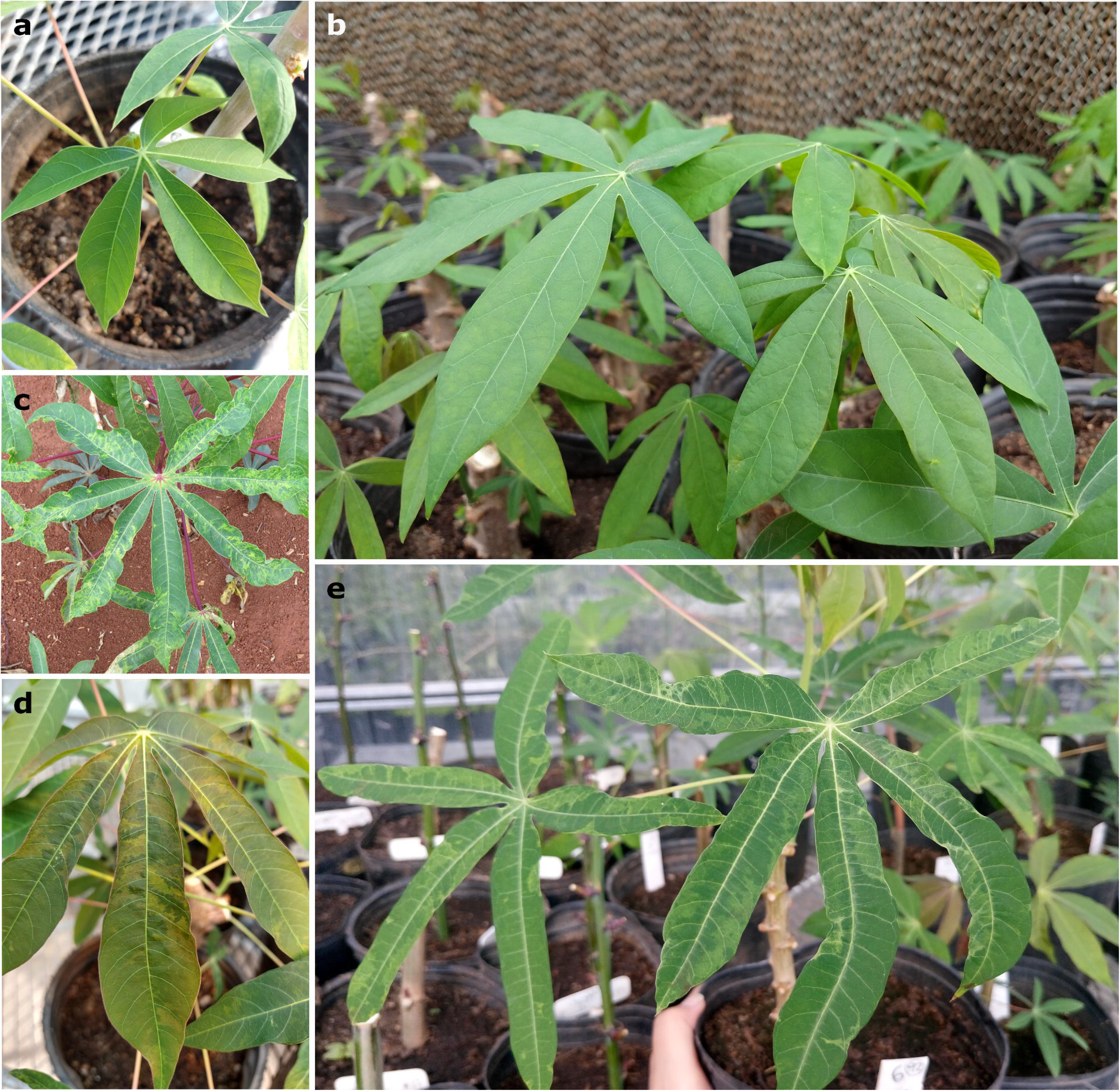
Plant material. (a, b) Healthy cassava plants (control). (c) Argentine field-collected cassava plant showing cassava common mosaic disease symptoms. (d, e) CsCMV-infected cassava plants 90 days after planting (DAP), showing mosaic leaf symptoms with alternating green and yellow patches. Representative plants were used for pictures.

The structure of chloroplasts revealed abnormalities in mesophyll CsCMV-infected cells, which included dramatic chloroplast malformations, such as small vesicles or vacuoles in stroma, disorganized grana stacks and irregular arrangement of stroma lamellae (Fig. 2). Furthermore, CsCMV, as a potexvirus member, have flexuous filamentous particles of 470–580 nm in length, which were observed accommodated in inclusion bodies of fibrous masses containing these filamentous particles, running more or less parallel to each other, but not in a definite arrangement, in the cellular cytoplasm near the chloroplast membrane (Fig. 2).

**Fig. 2.**
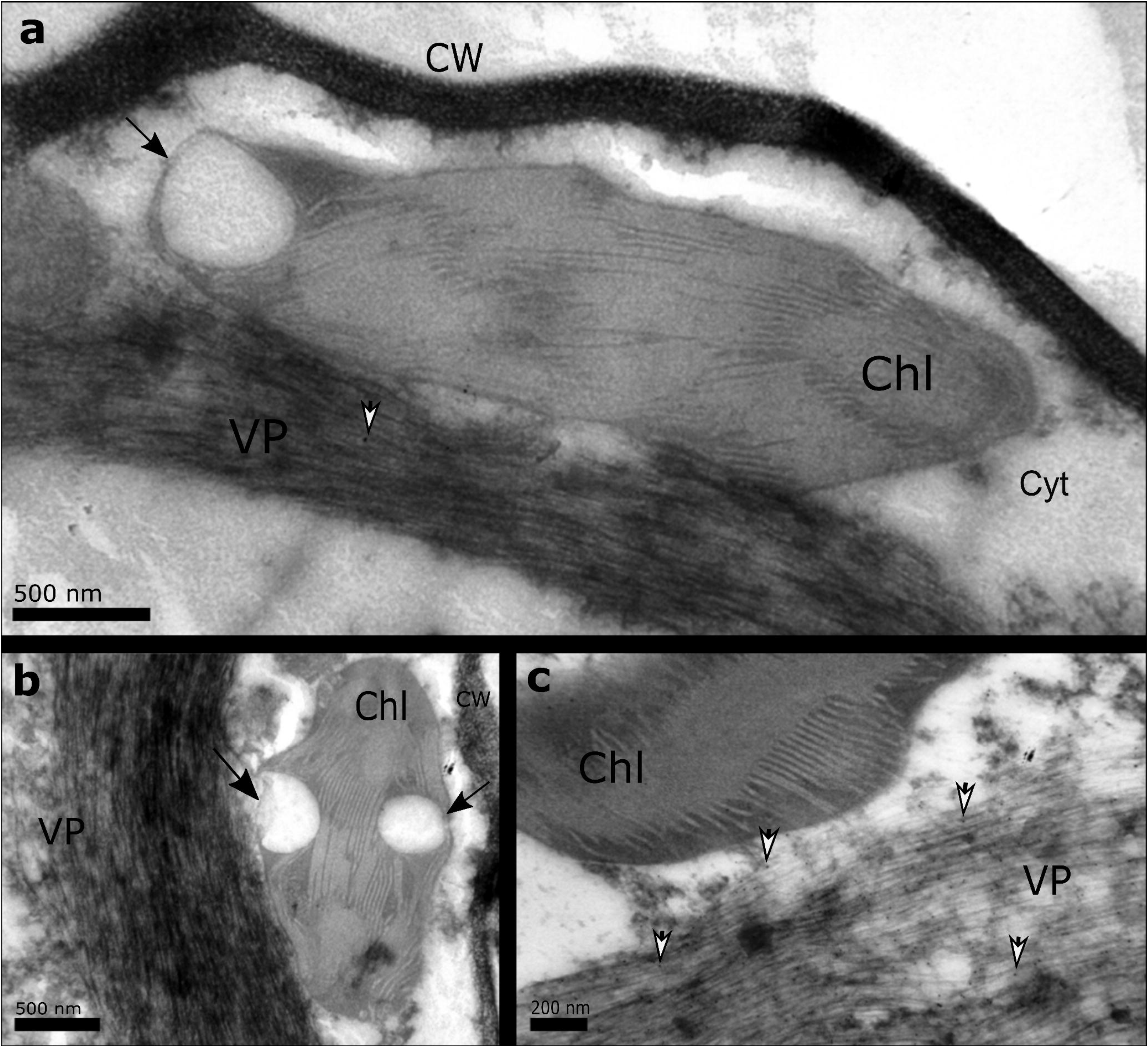
Ultrastructure of chloroplasts from CsCMV-infected mesophyll cassava cells. Transmission electron micrographs of a small sample (0.2 cm wide × 0.5 cm long) slice from the middle part of leaf tissues from CsCMV-infected cassava plants. Leaf samples were prepared for TEM and the capsid protein immunogold labeling of CsCMV was performed as described in material and methods. (a, c) Capsid protein gold particle deposits (white head arrows) were located in cytoplasm around chloroplasts (Chl), where virus particles (VP) were present. CW-cell wall, Cyt-cytoplasm. (a, b) Black arrows indicated chloroplast malformations, such as small vesicles or vacuoles in stroma. Bars = 500 nm (a, b) and 200 nm (c)

### Photosynthetic activity is compromised in CsCMV-infected leaves

Fast ChlF kinetics was measured to reflect the decrease of PSII activity induced by CsCMV in cassava plants. The value of the OJIP parameters, affected by the CsCMV, were normalized to the control ones and plotted in a radar graph to deploy a global view of the structure and function of PSII electron transport (Fig. 3b). Accordingly, the maximum quantum efficiency F_v_/F_m_ was negatively affected as the disease progressed, becoming more evident after 90 DAP. Such changes occurred due to the reduction of both minimum (F_o_) and maximum (F_m_) ChlF emission. The energy flow taking place at the reaction centers (RC) was also altered by the CsCMV infection. Such event begins at light absorption (ABS) by PSII antenna pigments and ends at reduction of the end electron acceptors at the photosystem I (PSI) electron acceptor side (RE) driven by PSI. As of 60 DAP, a continuous energy loss as latent heat (DI_o_/RC) was observed. Then, changes in the size of the antenna complex (ABS_o_/RC) and energy capture (TR_o_/RC) took place after 75 DAP. PSI electron acceptor side (RE_o_/RC) also increased during the same period. Absolute performance index (PI_abs_) of infected plants slumped by 33% after 60 DAP with respect to non-infected plants. Likewise, total performance index (PI_total_) diminished, but recovered slightly 90 DAP. Significant symptoms of chlorosis of cassava leaves (SPAD index; Fig. 3b) were evident as early as the 60 DAP. As a consequence of chloroplast damage and the energetic imbalance at the PSII electron transport, reducing power for photosynthesis was reduced. This result may explain the reduction of the photosynthetic rate observed 90 and 210 DAP (13.5% to 24.2%, respectively) (Fig. 3a).

**Fig. 3.**
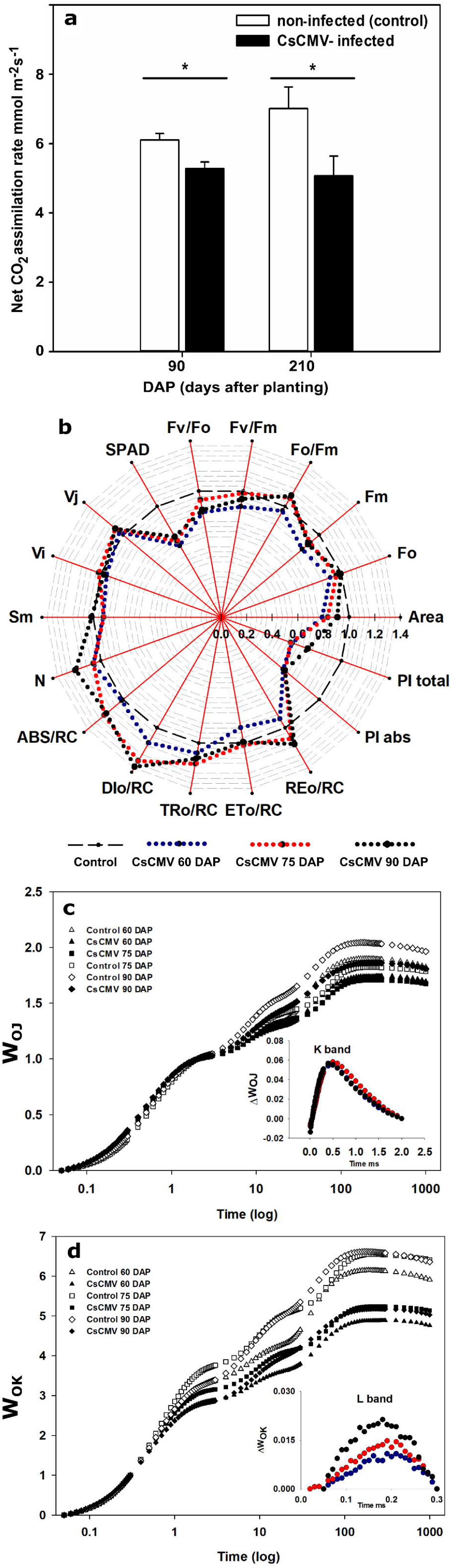
PSII activity and CO_2_ fixation in CsCMV-infected leaves. (a) Photosynthetic rates of non-infected (control) and CsCMV-infected cassava plants at two growth stages, 90 and 210 days after planting (DAP). (b) Radar plot showing OJIP test parameters in cassava control plants and CsCMV-infected. Measurements were carried out at 60 (blue), 75 (red) and 90 (black) DAP. Parameter abbreviations are described in Supplementary Table S1. (c) Changes in the oxygen evolving complex (OEC, K band) and (d) energetic connectivity of PSII units (L band) in non-infected (empty icons) and CsCMV-infected cassava plants (black circles). (c, inset panel) WOJ: Normalized fluorescence kinetics from O (50 μs) to J step (2 ms). ΔWOJ =WOJ_(Treated)_ -WOJ_(Control)_, kinetics difference between treated and control plants, revealing the K band. (d, inset panel) WOK: Normalized fluorescence kinetics from O (50 μs) to K step (300 μs). ΔWOK =WOK_(Treated)_ -WOK_(Control)_; kinetics difference between treated and control plants, revealing the L-band at 60 (blue), 75 (red) and 90 (black) DAP

The differences of the normalized ChlF curves (ΔW_OK_ and ΔW_OI_) revealed the appearance of hidden steps, such as the K and L bands (OLKJIP). Positive bands (K band) occurring at about 300 μs are considered to denote failure at the OEC during 60, 75, and 90 DAP in CsCMV-infected plants (inset Fig. 3c; Oukarroum et al 2007). Moreover, the fluorescence rise during the first 150 μs (L-band) is attributed to the loss of energetic connectivity between PSII units (Ripoll et al 2016). Here, the appearance of L band increased from 60 to 90 DAP (inset Fig. 3d).

### CsCMV infection induce redox imbalance in cassava plants

The FRAP assay offers a putative index of antioxidant or reducing state. An indirect estimation of the total non-enzymatic antioxidant content of the extract showed significant differences between the means of CsCMV-infected plants and the control plants (Fig. 4). The reduction of the antioxidant capacity was about 20.8% in virus-infected plants.

**Fig. 4.**
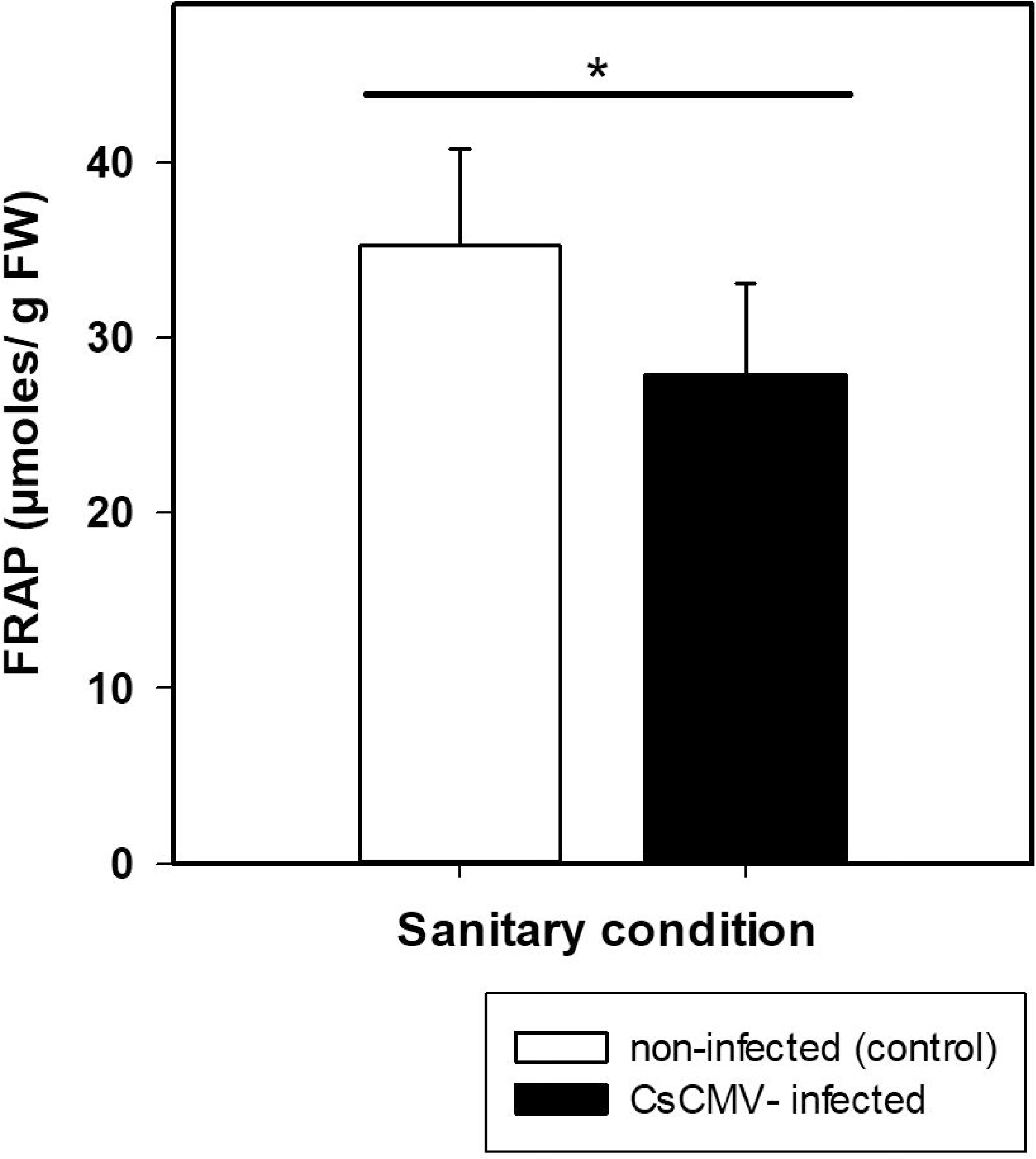
Antioxidant system in CsCMV-infected leaves. FRAP was measured in cassava leaves at 90 DAP. Means of non-infected (white bars) and CsCMV-infected (black bars) plants. FW: fresh weight * statistically significant according to ANOVA followed by LSD Fisher test (p < 0.05) (n = 30)

### CsCMV infection: effects on carbohydrate and starch accumulation

Starch content in infected cassava leaves decreased significantly (by 22.8%), with respect to control (Fig. 5a). Total soluble sugar content (Fig. 5b) and AGPase activity were not affected in infected leaves (Fig. 5c).

**Fig. 5.**
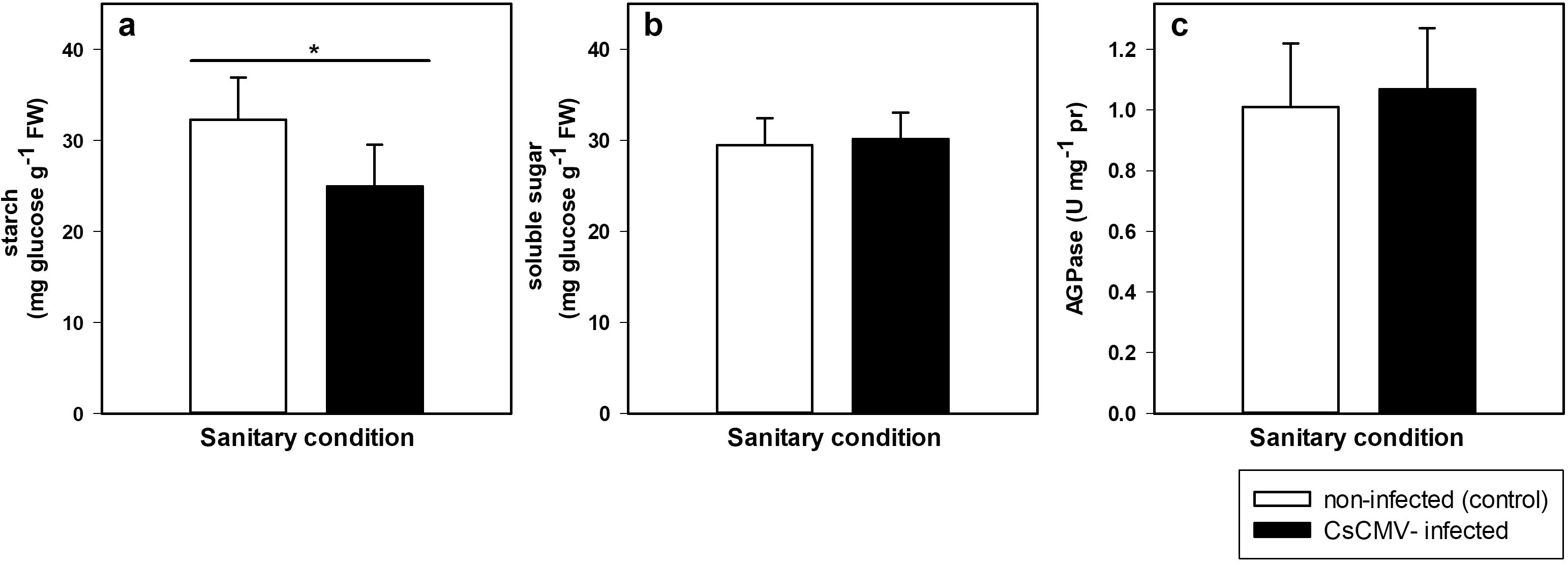
Carbon metabolism on cassava-CsCMV interaction at 90 DAP. (a) Starch and (b) total soluble sugar content (n = 30 for each sanitary condition). (c) AGPase activity of cassava leaves of two sanitary conditions (n = 40). FW: fresh weight; pr: protein * statistically significant according to ANOVA followed by LSD Fisher test (p < 0.05)

A highly detailed profile of soluble sugars evidenced a decrease in maltose content without significant differences in fructose, glucose or sucrose in CsCMV-infected leaves with respect to control plants (Table 1). The sucrose/starch ratio showed a significant increase with respect to non-infected plants (Table 1).

**Table 1.**
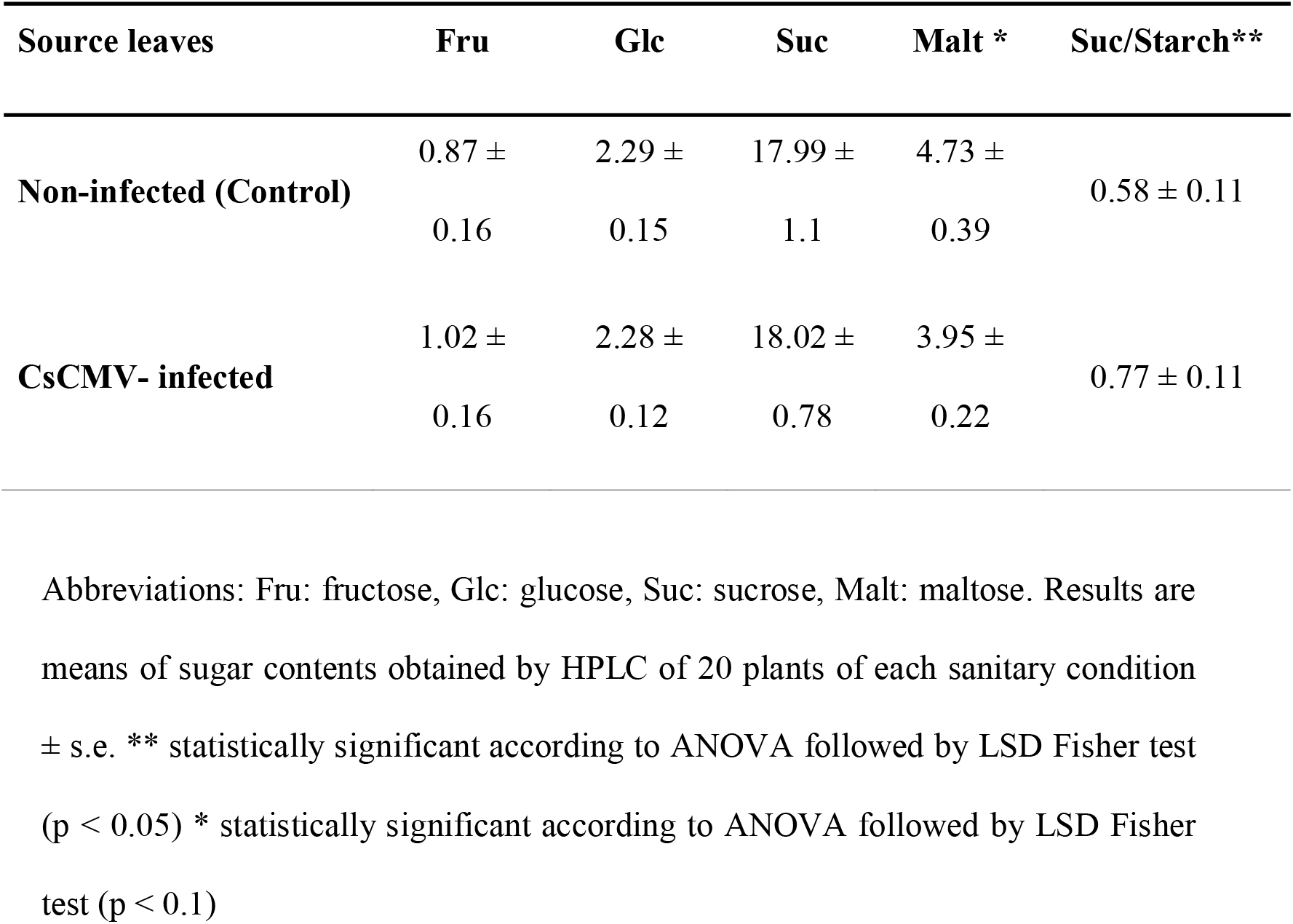
Sugar profile (mg g^−1^ FW) on CsCMV-infected leaves at 90 DAP. Abbreviations: Fru: fructose, Glc: glucose, Suc: sucrose, Malt: maltose. Results are means of sugar contents obtained by HPLC of 20 plants of each sanitary condition ± s.e. ** statistically significant according to ANOVA followed by LSD Fisher test (p < 0.05) * statistically significant according to ANOVA followed by LSD Fisher test (p < 0.1).

## Discussion

Cassava-CsCMV compatible viral infection causes 30-60% yield reduction of cassava roots (Costa and Kitajima 1972; Venturini el al. 2016; Zanini et al. 2020). These yield losses reflect the final result of a series of physiological events that occurred at earlier stages of viral interaction. The present study was focused on chloroplast morphological alteration and functional effects on source leaves during the course of viral infection. The mosaic symptoms induced by CsCMV were associated with structural changes of chloroplasts, decrease in relative Chl content, electron transport rate alteration, photoinhibition, reduction of CO_2_ fixation, reduction of starch, and maltose accumulation associated with increase in sucrose/starch ratio and drop of total antioxidant content.

The interaction between chloroplast and the invading virus plays a critical role in viral infection and pathogenesis (Zhao et al. 2016). Our previous results showed inclusion bodies of fibrous masses containing filamentous particles in the cellular cytoplasm near the chloroplast membrane (Zanini et al. 2014). In the present study, which was focused on chloroplast structure, we observed extrusion of the chloroplast membrane with amoeboid-shaped appearance, among other results (Fig. 2b). Several research works have shown that viruses can interact with chloroplast proteins and induce the formation of membrane vesicles during viral replication, which impairs chloroplast functions in plants (Liu et al. 2014; Li et al. 2016a; Zhao et al. 2016; Souza and Carvalho 2019). For example, high levels of *Potato virus X* (PVX, *Potexvirus*)-Coat Protein (CP) caused structural alteration of the chloroplast membranes, thylakoid grana and invagination of the cytoplasm within these organelles; it was also found to interact with the plastocyanin (Zhao et al. 2016).

The development of mosaic symptom induced by CsCMV not only was a consequence of the chloroplast structure alteration in cassava mesophyll cells (Fig. 2), but also severely inhibited its function (Fig. 3). Our results agree with findings of Liu et al. (2014), who reported that visible symptoms of chlorosis in cassava plants infected with *African cassava mosaic virus* (ACMV, *Begomovirus*) were associated with chlorophyll breakdown (reduction of chlorophyll content and fewer grana stacks) and upregulation of chlorophyll degradation transcript genes. For a long time, the chloroplast-virus interaction has been a matter of debate due to its complexity. We noticed that CsCMV infection negatively affected the CO_2_ gas-exchange rate, with a constant fall over the progress of the disease (Fig. 3a). This phenomenon could be explained by the lower production of ATP and NADPH, due to the closure of RC, as indicated by the increment in ABS_o_/RC values (Fig. 3b). On the other hand, virus infection could have reduced the carboxylation capacity through impairment of Rubisco activity (Sampol et al. 2003). According to the analysis of ChlF signals, CsCMV showed a severe impact on the photosynthetic light primary reaction at 90 DAP (Fig. 3b).

The loss of PSII electron transport efficiency is regarded as a common phenomenon under biotic stress. As expected, the OJIP-test showed significant changes of most of the ChlF parameters over time in CsCMV infected plants in relation to the healthy ones. For instance, the viral infection decreased the photochemical efficiency of PSII (F_v_/F_m_; Fig. 3b), whereas it increased light energy dissipation (DI_o_/RC) significantly, and decreased performance index (PI_abs_) and alteration in the linear electron transport (ET_o_/RC) in PSII. According to Gallé and Flexas (2010), a decrease in F_v_/F_m_ is due to an increment in thermal dissipation (non-photochemical quenching) at the expense of photochemical activity. Furthermore, when this parameter becomes markedly reduced, it is considered as an indication of photoinhibition or PSII photo-damage (Maxwell and Johnson, 2000), as is the case of CsCMV infected-leaves at 90 DAP. Similar results were found in *Sunflower chlorotic mottle virus* (SuCMoV, *Potyvirus*), with degradation of D1 protein, an essential protein of PSII functionality, resulting in photoinhibition/photodamage, and drop of F_v_/F_m_ parameter (Rodríguez et al. 2010).

Oxygen evolving complex (OEC), a Mn_4_O_5_Ca protein cluster embedded in the PSII, is essential in the photosynthesis process. It has been demonstrated that the viral CP, or some other products synthesized as a result of the viral infection, decreases the amount of OEC polypeptides, as reported in tobacco leaves infected with *Cucumber mosaic virus* strain Y (CMV(Y), *Cucumovirus*) (Takahashi and Ehara 1992). In our work, a progressive loss of OEC activity was revealed by the K-band starting 60 DAP in CsCMV-infected leaves (inset Fig 3c). Under particular stressful conditions, the K-band was demonstrated to indicate a disruption of the OEC (Kalaji et al., 2016). Likewise, the OJIP analysis evidenced that the slope between F_o_ and F_150_ (L-band) was sensitive to the CsCMV attack, which was progressively increased up to 90 DAP (inset Fig. 3d). This phenomenon occurs due to a loss of “connectivity” within the tripartite system (core antenna-LHCII-Reaction Centre) under particular abiotic stress conditions (Strasser et al. 2004; Kalaji et al., 2016); here it is reported for virus infection for first time.

The virus ability to impair chloroplast function and disrupt the photosynthetic electron transport chain ultimately leads not only to the decrease of the carboxylation activity but also to ROS increase (Rodríguez et al. 2010; 2012; Souza et al. 2019). CsCMV induced redox alteration demonstrated by the lower antioxidant capacity (FRAP), which might indicate an oxidative stress process. Furthermore, the CO_2_ fixation decrease might be directly related to soluble sugar production. Nuwamanya et al. (2017) demonstrated that cassava brown streak disease (CBSD), caused by potyviruses, leads to a general reduction in primary carbohydrate composition (total reducing sugar and starch contents) in leaves; these effects correlated with the negative effect on photosynthetic apparatus of infected leaves. However, our results did not show such significant differences in the total soluble sugar contents in CsCMV-infected source leaves with respect to non-infected plants. The differences between results might be related to the specific source tissue analyzed during our work. These results led us to ask about the origin of these sugars, since photosynthetic activity was decreased by viral infection. One possible answer might be related to the findings of Sharkey et al. (1985) that, at low rates of photosynthesis, carbon would preferably be used for sucrose synthesis rather than for storage in starch form. Accordingly, present study showed a rise of the ratio of carbon partitioned into sucrose versus starch (suc/starch ratio, Table 1) in CsCMV-infected plants. Another possible answer may be starch breakdown, which provides a source of reduced carbon when photosynthesis cannot occur. Indeed, starch content was decreased during viral infection (Fig. 5a). However, maltose, which might be increased in leaves when starch catabolism is induced, showed a significant decrease in CsCMV-infected leaves (Table 1). In this context, we cannot rule out the fast maltose metabolism to viral replication. Moreover, the different sugars might be finely sensed and might provide a mechanism to conduct carbon flux during viral infection (Sade et al., 2013; Hulsmans et al. 2016). Besides metabolic flux diversion, another possible source of soluble sugar could be related to alteration of sugar transporters, which convert these `source` leaves into `sink` tissues (Kanwar and Jha 2019), or a gluconeogenesis process induction. These possible sugar sources are not mutually exclusive but might be operating in combination, and have to be explored in future studies.

To conclude, cassava produces storage roots, which are the major sink for storing starch derived from carbohydrate partitioning and long-distance sugar transport from the leaves (Yan et al. 2019). The early alterations in photosynthesis and chloroplast structure observed during the present work might be related to negative impact on yield previously observed at a later stage (Zanini et al., 2020). Such changes imply smaller reserve roots, i.e., fewer accumulated carbohydrates. Therefore, understanding the metabolic alteration induced by CsCMV in source leaves at this early stages is critical for improving cassava yield. This is the first report demonstrating physiological alterations associated with chloroplast ultrastructure and function in the cassava-CsCMV pathosystem.

## Abbreviations

AGPase: ADP-glucose pyrophosphorylase
ChlF: Chlorophyll-a fluorescence
CsCMV: *Cassava common mosaic virus*
DAP: Days After Planting
FW: Fresh Weight
OEC: Oxygen Evolving Complex
PSII: Photosystem II
RC: Reaction Center
ROS: Reactive Oxygen Species

## Author Contribution

MR and AAZ designed the experiments. AAZ conducted the experiments. AAZ and MR took the measurements. AC carried out in vitro culture and acclimatization of cassava seedlings. LDF contributed with cassava plants and virus detection. DFL contributed in the fluorescence analysis. AAZ and PP performed the statistical analysis of data. AAZ and MR drafted and co-wrote the manuscript. AAZ, LDF, DFL and MR contributed with the discussion and editing of the manuscript. MR conceived and supervised the project.

## Acknowledgements

We are grateful to Claudia Nome and Leandro Ortega for helpful assistance in electron microscopy and HPLC techniques, respectively. We thank Edith Taleisnik for comments on the manuscript and helpful discussions. This work was supported by grants from the Agencia de Promoción Científica y Tecnológica, Argentina (PICT-2014-551), and the Instituto Nacional de Tecnología Agropecuaria (INTA-I085). AAZ is fellow and MR is researcher of CONICET (Consejo Nacional de Investigaciones Científicas y Técnicas, Argentina). MR and LDF are researchers of INTA.

