## Supplementary data for "*Cassava common mosaic virus* causes photosynthetic alterations associated with changes in chloroplast ultrastructure and carbohydrate metabolism of cassava plants"

Affiliation and address:

^1^ Instituto de Fisiología y Recursos Genéticos Vegetales, Centro de Investigaciones Agropecuarias-INTA, Av. 11 de Septiembre 4755, X5020ICA, Córdoba, Argentina.

^2^ Unidad de Estudios Agropecuarios (UDEA- CONICET)

^3^ Instituto de Patología Vegetal, Centro de Investigaciones Agropecuarias-INTA

^4^ Unidad de Fitopatología y Modernización Agrícola (UFYMA- CONICET)

^5^ Cátedra de Estadística y Biometría de la Facultad de Ciencias Agropecuarias de la Universidad Nacional de Córdoba, grupo vinculado a UFYMA- CONICET

^6^ Centro de Investigación y Transferencia de Formosa - CONICET. Instituto Universitario de Formosa, Facultad de la Producción y del Medioambiente, UNaF

Av. 11 de Septiembre 4755,

X5020ICA, Córdoba, Argentina.

**List of Supplementary Figures:**

**Fig. S1:** Leaf area.

**Fig. S2:** Cassava plant elongation rates at four growth stages.

**Fig. S3:** Number of nodes and leaves per bud.

**Fig. S4**: Ultrastructure of chloroplasts from non-infected mesophyll cassava cells.

**Table S1**: Formulae and glossary of terms used by the JIP- test

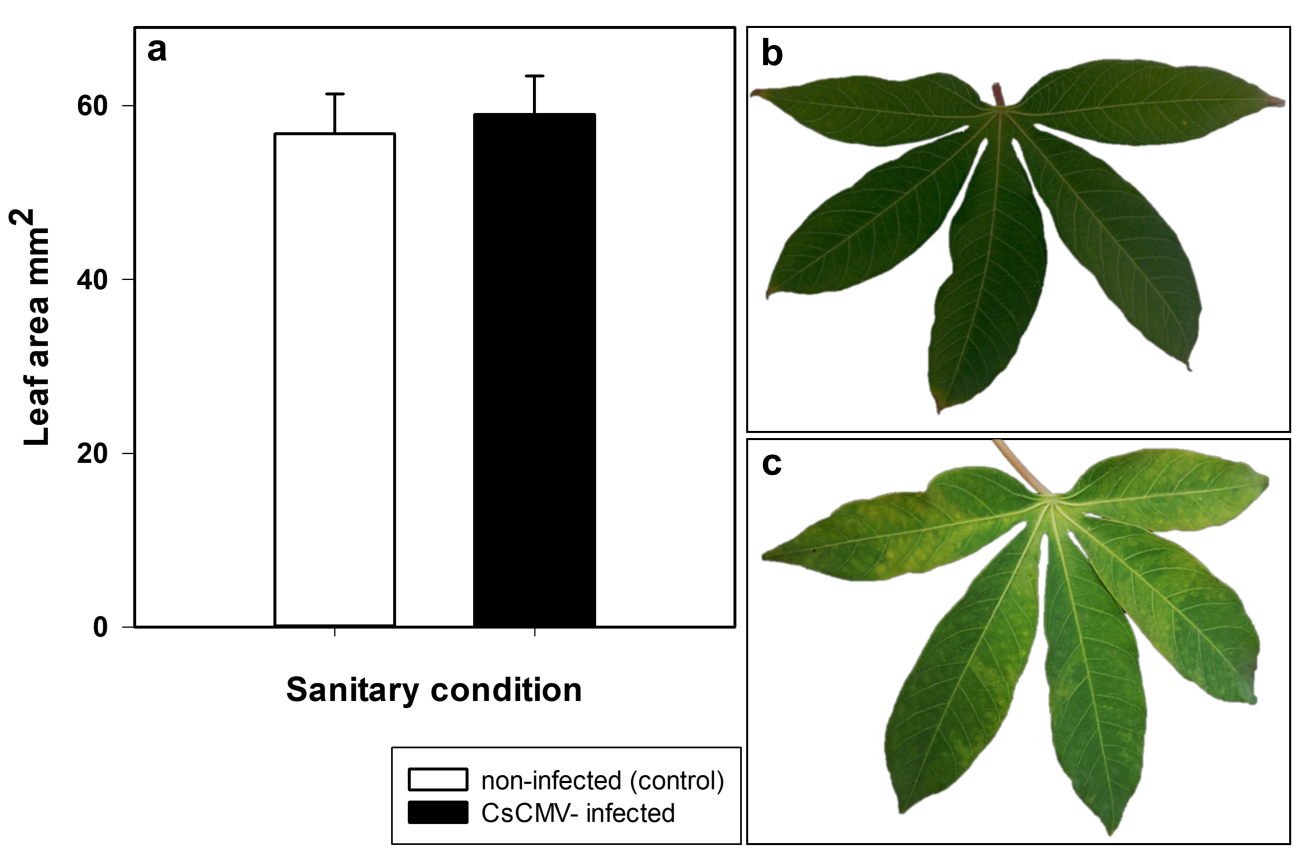

**Fig. S1:** Leaf area. (a) Means obtained of the leaf area for each sanitary condition at 90 DAP (N = 60). There are no significant differences by LSD Fisher test. Morphology of the fourth leaf (from the bottom up) of (b) non-infected (control) and (c) CsCMV- infected cassava plants, at 90 DAP.

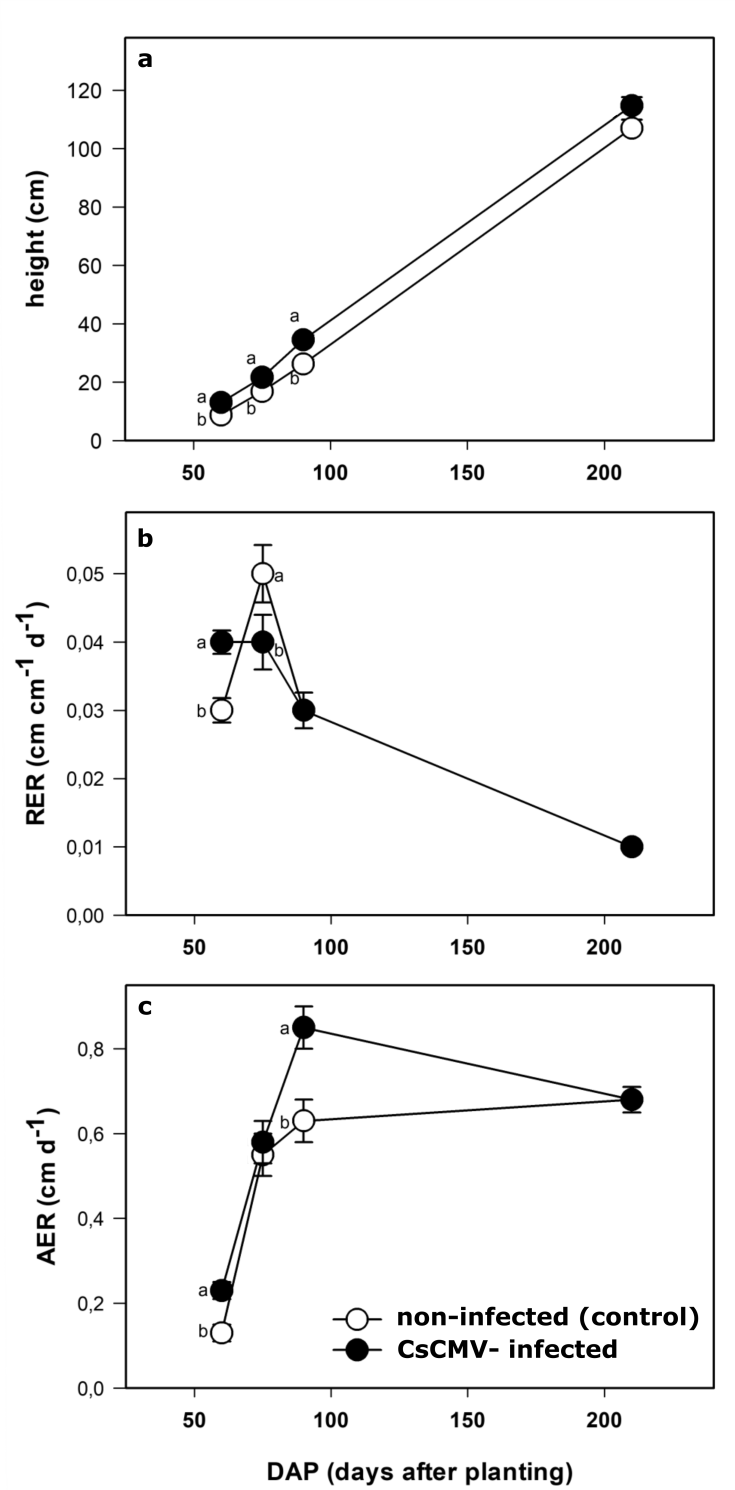

**Fig. S2:** Cassava plant elongation rates at four growth stages. (a) Shoot height at four growth stage (60, 75, 90 and 210 DAP). (b) RER- relative elongation rate and (c) AER- absolute elongation rate of cassava plants. Different letters indicate statistically significant differences (p < 0.05) by LSD Fisher (n = 30). AER (cm day^-1^) was defined as the increase in height per unit of time and the RER (cm cm^-1^ day^-1^) as the increase in height per unit of height of the existing plant and per unit of time. The equations were defined as: $AER= (A_{2}-A_{1})/(t_{2}-t_{1})$; $RER= ({ln A}_{2}-{ln A}_{1})/(t_{2}-t_{1})$. Where A_2_ and A_1_ are the height of the plant at times 2 (*t*_2_) and 1 (*t*_1_)

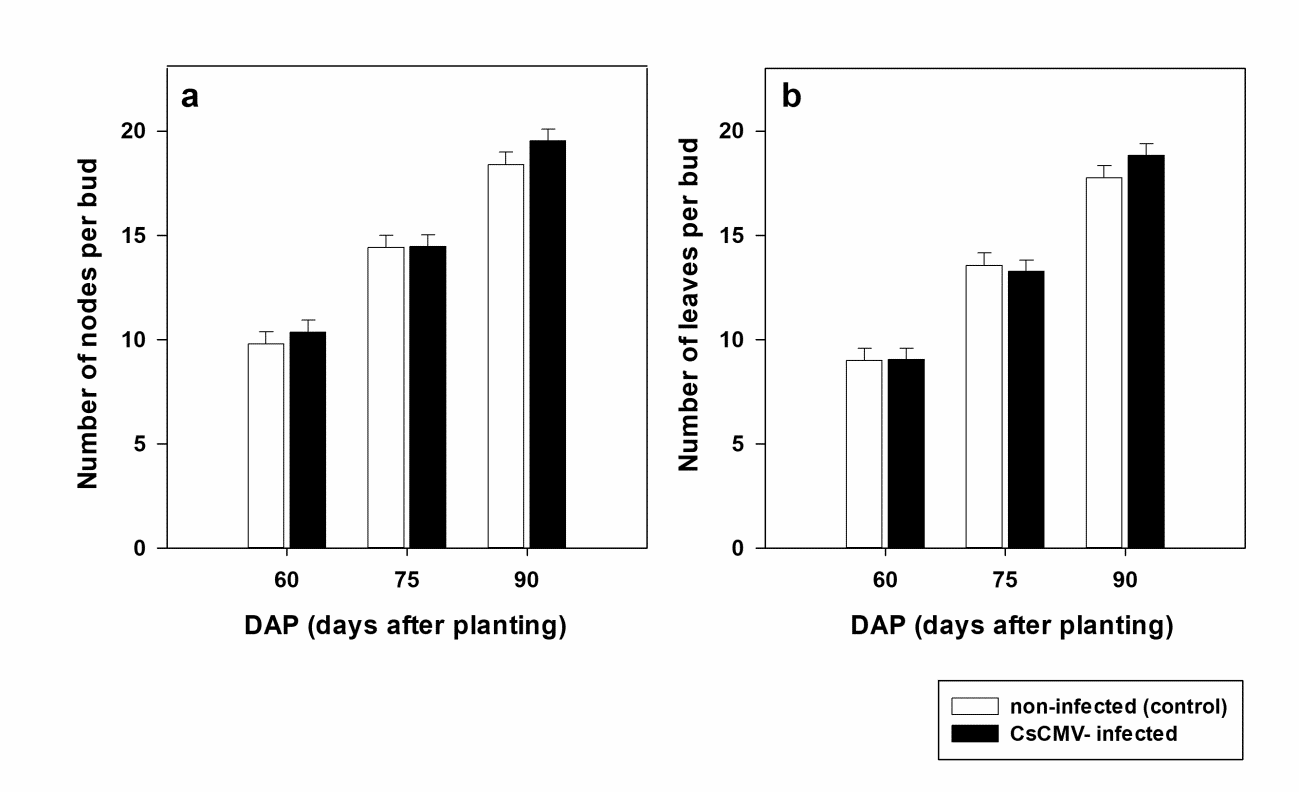

**Fig. S3:** Number of nodes and leaves per bud. (a) Number of nodes and (b) leaves per bud. Counting at three growth stages of cassava plants without (control) and with virus infection. No significant differences were observed by LSD Fisher test

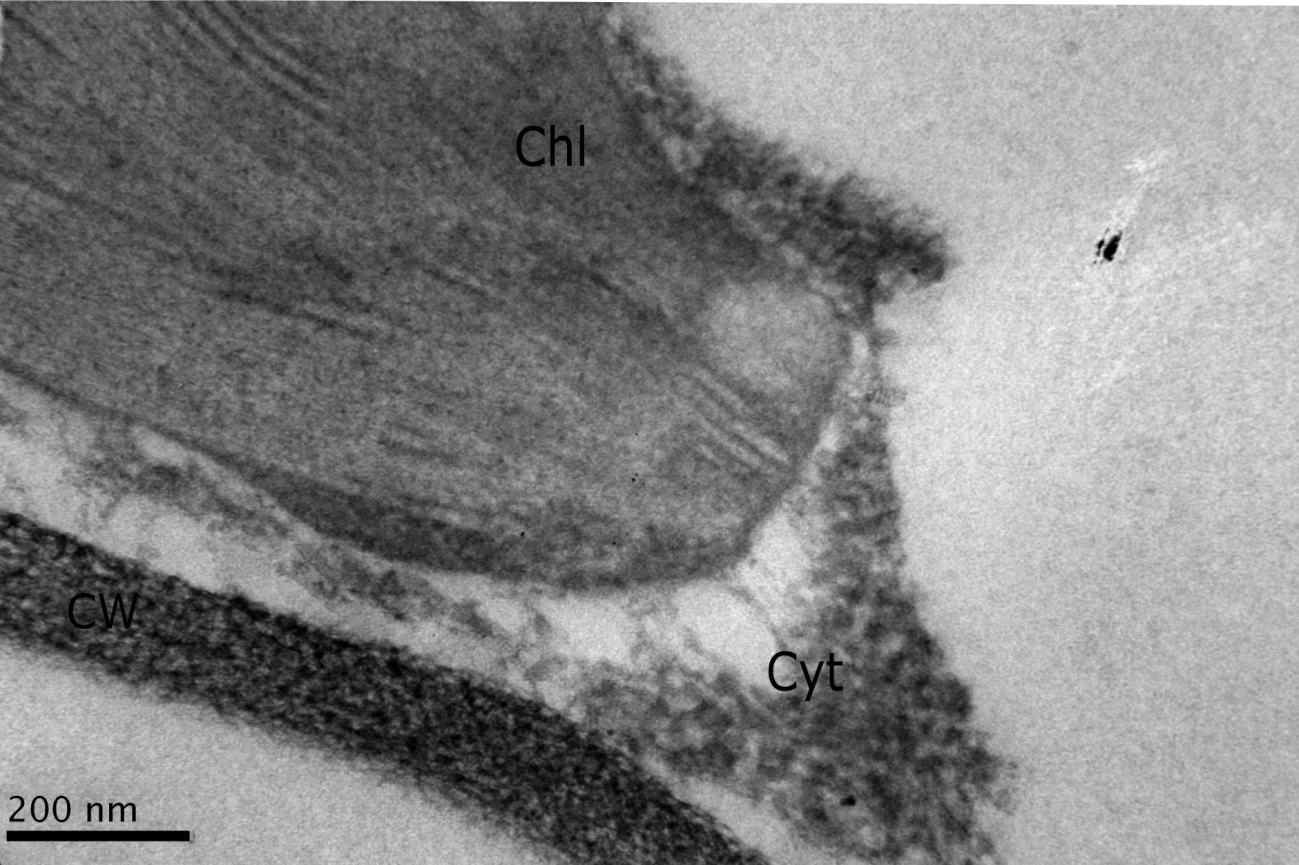

**Fig. S4**: Ultrastructure of chloroplasts from non-infected mesophyll cassava cells. Transmission electron micrographs of a small sample (0.2 cm wide × 0.5 cm long) slice from the middle part of leaf tissues from virus-free cassava plants. Leaf samples were prepared for TEM and the capsid protein immunogold labeling of CsCMV was performed as described in material and methods. Chl- chloroplast, CW- cell wall, Cyt- cytoplasm. Bars = 200 nm

**Supplementary Table S1**. Formulae and glossary of terms used by the JIP-test (modified from Strasser et al. 2004)

| **F_o_** | Estimated emission by excited chlorophyll a in PSII reaction centers (RCs) after dark adaptation, when the first stable electron acceptor in PSII (Q_A_) is fully oxidized. |
| --- | --- |
| **F_m_** | Maximum fluorescence obtained at saturating light intensity and when Q_A_ is fully reduced. |
| **F_v_** | Maximum capacity for photochemical quenching. Calculated as F_v_=F_m_-F_o_ |
| **F_v_/F_m_** | Maximum quantum efficiency of PSII. Its values decrease in stressed plant samples. |
| **F_v_/F_o_** | Size and number of active reaction centers RC. Reflect impairment and downregulation of PSII photochemistry and low electron transport. |
| **V_j_** | Relative variable fluorescence at 2 ms, step J. Calculated as V_j_=(F_J_ – F_0_)/(F_m_ – F_0_). |
| **V_i_** | Relative variable fluorescence at 30 ms, step I. Calculated as V_i_=(F_i_ – F_0_)/(F_m_ – F_0_). |
| **Sm** | Measure of the energy needed to close all reaction centers. |
| **N** | Q_A_ turnover number, it indicates how many times Q_A_ has been reduced in the time span from 0 to t_Fmax_ . |
| **Area** | Total complementary area between fluorescence induction curve and F = F_m_ |
| **Dl_o_/RC** | Ratio of total dissipation to the amount of active RCs. It increases due to the high dissipation of the inactive RCs. |
| **TR_o_ / RC** | Trapped energy flux per RC (at t=0). |
| **ER_o_ / RC** | Electron transport flux per RC (at t=0) |
| **RE_o_ / RC** | Electron flux reducing end electron acceptors at the PSI acceptor side, per RC |
| **PI ABS** | Performance index on absorption basis, it is an indicator of sample vitality. ABS is flux of photons absorbed by the antenna pigments Chl |
